# EcoXAI: Autonomous Agentic Ecosystem for Explainable Artificial Intelligence and Biomedical Discovery

**DOI:** 10.64898/2026.07.08.737358

**Authors:** Nicholas Matsumoto, Hyunjun Choi, Philip J. Freda, Alena Orlenko, Mythreye Venkatesan, Miguel E. Hernandez, Zhiping Paul Wang, Jason H. Moore

## Abstract

**Motivation:** As biomedical datasets and knowledge graphs continue to grow in size, complexity, and heterogeneity, navigating and extracting actionable insights from them presents a major bottleneck for researchers. There is a clear need for autonomous analytical solutions that can utilize recent advancements in agentic AI such as agent harnessing and loop engineering without introducing hallucination or workflow fragmentation. Researchers, regardless of technical expertise, need tools that streamline complex data analysis and deliver meaningful, actionable insights grounded in both data and established biomedical knowledge. EcoXAI addresses this by introducing a modular, customizable, containerized multi-agent system that structures analysis into explicit pipeline execution stages, lowering the computational barrier for clinical and translational researchers.

**Result:** EcoXAI replaces monolithic AI text interfaces with an autonomous execution-driven framework with specialized bioinformatics agents for delivering proactive, data-driven insights grounded in established biological knowledge. Unlike purely LLM-driven or less integrated AI solutions prone to hallucinations or biologically implausible outcomes, EcoXAI’s multi-agent framework, which leverages modern agentic management and explicit knowledge graph integration, provides greater transparency and verifiability in its reasoning. In our use case in drug repurposing for Alzheimer’s Disease, EcoXAI evaluated 103 drug candidates and identified 79 novel candidates whose predictive models exceeded a randomized baseline, including the CCR5 antagonist Maraviroc, whose generated hypothesis was subsequently supported by the literature. These results demonstrate the potential of knowledge graph-grounded AI agents to accelerate hypothesis-driven biomedical research.

**Availability and implementation:** EcoXAI is available on GitHub at: https://github.com/EpistasisLab/EcoXAI.

## 1 Introduction

The increasing scale and complexity of biomedical data have created a need for analysis frameworks that can combine statistical rigor, biological prior knowledge, and flexible workflow execution. In biomedical data science, important questions often span multiple modes of evidence, including association testing, predictive modeling, predictive analytics, pathway interpretation, and literature or knowledge-graph support. However, conventional analytical pipelines typically address these tasks separately, while general-purpose AI systems may generate plausible but biologically ungrounded outputs. This disconnect is especially limiting in complex disease domains such as Alzheimer’s disease, where meaningful discovery requires both dataset-specific evidence and mechanistic biological context.

To address this challenge, we present an autonomous agentic Ecosystem for Explainable AI (EcoXAI), a knowledge graph-grounded multi-agent framework for iterative biomedical discovery. EcoXAI integrates structured biological knowledge with exploratory data analysis, hypothesis generation, and hypothesis-specific validation in a modular workflow. Rather than treating all analytical tasks as a single generic prediction problem, EcoXAI decomposes a user’s research objective into distinct stages: dataset normalization and profiling, domain knowledge retrieval from a user-provided comprehensive biomedical knowledge graph, generation of candidate hypotheses informed by both the dataset and the interaction with the knowledge graph, and validation using evaluation procedures appropriate to the hypothesis type. Supported hypotheses are retained in discovery memory, enabling subsequent rounds of analysis to build on prior results and avoid redundant hypothesis generation.

## 2 Implementation

EcoXAI is organized around discrete, auditable pipelines rather than a single monolithic agent. These pipelines include data ingestion, exploratory data analysis (EDA), data normalization, predictive analysis, knowledge-graph guided hypothesis generation, hypothesis execution, and memory management. The system interacts with large language models (LLM) through third party APIs or local deployments, and its modular design allows the orchestration layer to route tasks to specialized tools or agents as needed. Once the user uploads their dataset, EcoXAI’s EDA agent assesses and cleans the dataset and creates an explanatory analysis in a markdown file which will be included in every downstream agent’s memory. EcoXAI then performs the fully autonomous research loop: using the knowledge graph and prior experiments, if available, it provides a diverse set of hypotheses, then evaluates each hypothesis until the entire set is exhausted, and repeats this cycle until either token budget or number of hypotheses loops is reached.

EcoXAI’s architecture is built upon modern software engineering principles, employing a microservices approach facilitated by Docker containers. This containerized architecture ensures high portability and enables consistent, reproducible performance across diverse computing environments, providing a critical advantage in bioinformatics, where software deployment and dependencies often present challenges.

EcoXAI’s core intelligence relies on recent advances in AI harnessing frameworks, such as Claude Code, Codex, and OpenCode, which are software layers that equip an LLM with access to tools, specialized skills, and planning for complex workflows to perform real-world tasks reliably. These tools are particularly suited for biological data science problems requiring multi-step reasoning that need rather rigorous validation, creativity, and grounding. Agents are infused with customizable agent skills such as data normalization, exploratory data analysis (EDA), hypotheses generation/testing, and knowledge graph interaction.

A key feature is EcoXAI’s integration with public biological knowledge graphs, which grounds analyses in established biological knowledge within the graph, reducing the risk of hallucinations and implausible results, leading to enhanced interpretability by linking findings to known pathways and relationships as well as generating hypotheses with prior knowledge in context. This knowledge-aware approach is crucial for applications like genomic analysis or drug repurposing, where context is essential.

### 2.1 Key EcoXAI Features

1. **Multi-Agent Architecture and Workflow Orchestration**: EcoXAI employs a planner-orchestrator architecture that coordinates specialized agents and analytical services across the biomedical research workflow. Rather than relying on a single monolithic agent, tasks are delegated to components responsible for data ingestion and exploratory data analysis (EDA), knowledge graph interaction, hypothesis generation, predictive modeling, explainability, and reporting. The orchestrator determines execution order, manages concurrent hypothesis evaluations, and routes structured inputs and outputs between components. A diagram of the architecture is shown in Figure 1-1.
2. **Flexible Data Ingestion and Profiling**: EcoXAI supports multiple biomedical data formats and can ingest datasets containing demographic and clinical features, molecular features, and quantitative biomarkers, accommodating both classification and regression tasks. The EDA agent profiles feature distributions, missingness, class imbalance, correlations, outliers, and other quality or structure-related properties. It produces an explanatory Markdown report that is incorporated into the context of downstream agents, establishing dataset-specific grounding for subsequent reasoning.
3. **Knowledge Graph Grounding**: EcoXAI queries public biomedical knowledge graphs, such as the Alzheimer’s KnowledgeBase (AlzKB), using the Cypher query language to retrieve pathways, gene–disease associations, drug– target relationships, and other biological context. The knowledge graph agent automatically retrieves the schema of connected graph databases and constructs queries based on the available relationships. Retrieved information informs hypothesis generation, supports interpretation of model outputs, and helps prioritize biologically plausible candidates.
4. **Autonomous Hypothesis Generation**: EcoXAI generates candidate hypotheses by combining the user’s research question with dataset-derived signals and knowledge graph context. Each hypothesis specifies a candidate biological relationship or predictive signal, together with an analytical strategy for its evaluation. Prior hypotheses and their outcomes are retrieved from discovery memory through semantic similarity search, reducing redundant proposals and encouraging exploration of alternative biological relationships and feature-selection strategies.
5. **Parallelized Hypothesis Evaluation**: Each hypothesis is evaluated using a methodology suited to its type. Predictive hypotheses are assessed using machine learning performance metrics, while mechanistic or knowledge-graph-derived hypotheses are assessed using dataset-based corroboration and enrichment evidence. The execution agent translates each hypothesis into an executable analytical workflow, and modeling and explainability components train models, assess performance, and interpret relevant features. EcoXAI launches concurrent evaluation sessions according to user-defined settings and continues evaluating pending hypotheses until the current set is exhausted or an execution budget limit is reached. Upon completion, the orchestrator can initiate another hypothesis generation cycle.
6. **Discovery Memory and Auditable Execution**: EcoXAI maintains persistent records of prior hypotheses, validation outcomes, supporting evidence, execution scripts, and analytical reports. Hypotheses are stored as semantic vector embeddings, enabling Retrieval-Augmented Generation (RAG) and similarity search during subsequent generation cycles. Intermediate artifacts are shared with downstream agents through the orchestration layer, so later analyses build on previous findings rather than revisiting the same analytical space. Each pipeline stage keeps its own execution records, allowing individual experiments to be inspected, reproduced, or modified.
7. **Deployment and Modularity**: EcoXAI packages the analysis stack, including AI harnesses, specialized agents, and dataset management services, as Docker containers. This design promotes reproducibility and simplifies deployment across local, cloud, and collaborative environments. The system supports different LLMs, specialized tools, and user-defined agent skills, so researchers can adapt the workflow to available compute, task complexity, and institutional constraints. Individual agents and services can be modified or replaced without redesigning the framework.

**Fig 1.**
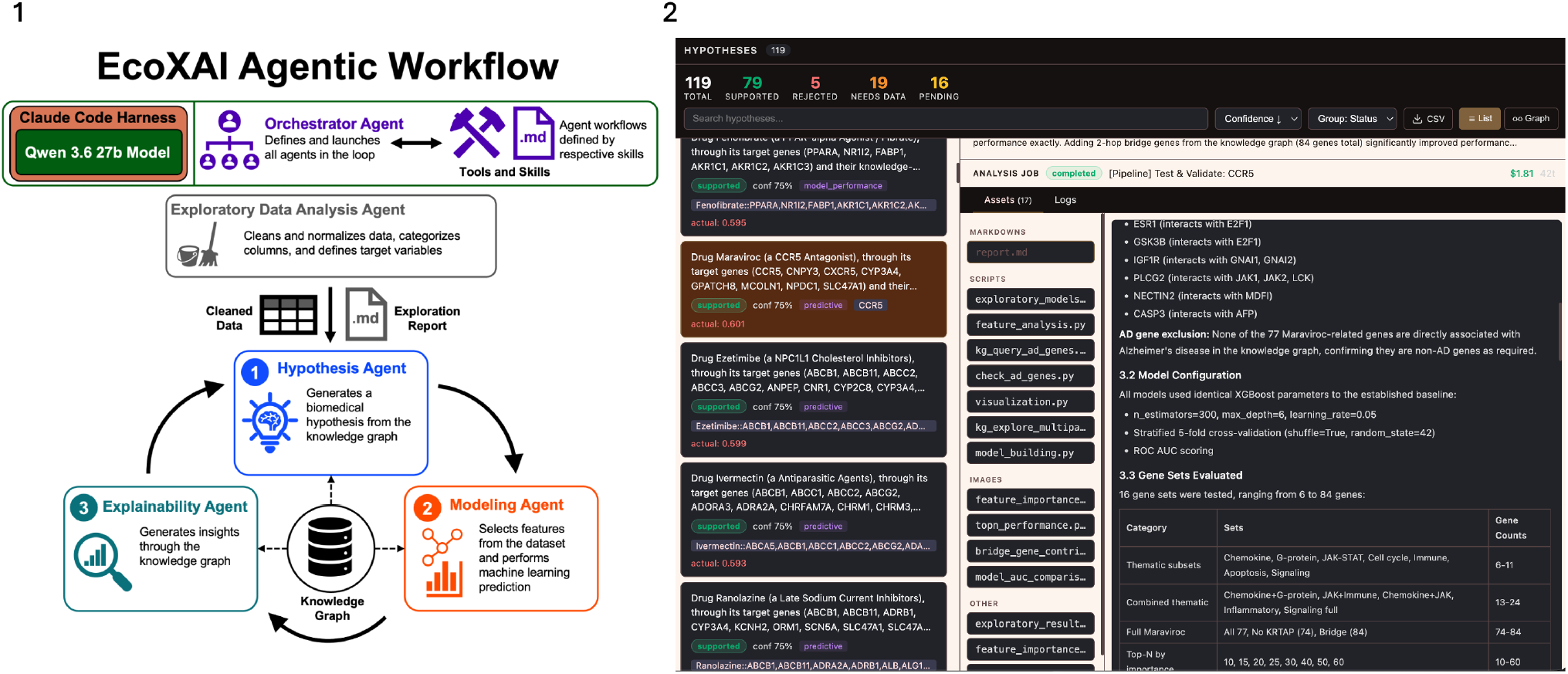
1) EcoXAI’s agentic workflow diagram. 2) Screenshot of the hypotheses page, which contain a list of all the generated hypotheses as well as the per hypothesis view. The hypothesis view contains assets that were generated from the evaluation of the hypothesis and a comprehensive report. 119 hypotheses were generated with 16 still pending execution. 79 of the 103 executed hypotheses have a cross-validated XGBoost AUC above the 0.5823 randomized baseline, which indicate potential drug candidates for Alzheimer’s disease.

## 3 Drug Repurposing Use Case

EcoXAI’s versatile capabilities support a wide range of bioinformatics applications and we will showcase a complex biomedical use case that display the full agentic properties of EcoXAI. To demonstrate EcoXAI’s ability to interact with grounded and transparent knowledge, we applied it to a publicly available Alzheimer’s Disease (AD) knowledge graph, AlzKB (https://AlzKB.ai/). AlzKB is a Memgraph graph database containing detailed data on genes, diseases, drugs, and other relevant entities, along with the semantic relationships between them (Romano et al., 2023). We use the 1.21 version of AlzKB for our benchmarks. EcoXAI’s skill management allows the agents to be capable of automatically retrieving the schema of any connected knowledge graph database, querying the requisite knowledge, and executing its task with it.

By interacting with AlzKB to analyze gene expression profiles or network contexts in AD, EcoXAI proposes and evaluates target candidates. In this case study, we tasked EcoXAI to not use drugs that are already associated with AD as to not use any genes that are connected to AD for its feature selection to force it to explore novel biological mechanisms. EcoXAI will generate machine learning models and test whether the AUC of the drug’s model is more performant in AUC than the randomized baseline.

### Gene scores

Genetic variants were obtained from the ADSP R4 v11 2023 release. Quality control removed rare variants (MAF *≤*1%), variants with low call rate (missing in *>*1% of samples), and poorly genotyped samples (missing *>*5%). Because individual variants typically have small effects sizes and often lack annotation, we aggregated the weighted effects of common variants within each gene into gene-level risk scores (GRS). This reduces hundreds of variants to one feature per gene and allows results to be interpreted at the gene level.

Variant effect sizes were taken from GWAS summary statistics conducted on held-out portion of the ADSP dataset after propensity score matching for population stratification (Orlenko et al., 2025). Variants were filtered for linkage disequilibrium using the PLINK2 clumping procedure (p1 = 1e-4, p2 = 1, r^2^ = 0.1), and those with GWAS p *≤* 0.05 were retained for gene mapping. Retained variants were annotated with the Variant Effect Predictor v113 and mapped to genes if located within the gene region. Gene start and end coordinates were taken from the Ensembl release 113 GTF annotation (GRCh38). For each individual *i* and gene *g*, the GRS was calculated with PLINK2 `--score` as

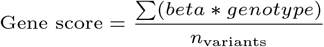

This yielded GRS for 20,935 genes in 11,280 ADSP individuals.

## 4 Results

Using a local Qwen 3.6 27B model running continuously for two days accruing approximately 50 million input tokens and 10 million output tokens, which would have otherwise cost over an estimated $300 using Anthropic’s Sonnet 4.6 token pricing, EcoXAI autonomously generated, executed, and evaluated 103 independent drug repurposing hypotheses. For each candidate drug, the framework interacted directly with the AlzKB knowledge graph to retrieve drug targets, identify biologically connected genes through multiple graph traversal strategies, construct competing feature sets, train XGBoost models with stratified 5-fold cross validation on the training set (80/20 split, random seed 25), and automatically generate a complete analytical report describing the resulting hypothesis.

Rather than optimizing a fixed feature set, EcoXAI iteratively explored multiple biological hypotheses for each drug. Through AlzKB and starting from experimentally validated drug targets, the framework autonomously generated alternative gene sets including direct targets, first- and second-order interactors, shared biological processes, pathway-restricted subnetworks, and other methods that the agent creatively decided to pursue without explicit human instruction. Each hypothesis was independently evaluated against a randomized baseline result of 0.582, which consisted of 1,000 XGBoost models trained on random gene sets containing up to 100 genes. For reference, there are 11 drugs in AlzKB that directly treat AD and each drug’s test AUC has exceeded the randomized baseline ranging from 0.584 to 0.665, which are modest in absolute terms, but they are the relevant comparison for drug candidates. We extended the gene inclusion criteria to two hops and up to 100 genes and found that the AD drug Selegiline achieved the top AUC of 0.665 with only 56 genes.

Of the 103 candidate drugs evaluated, 79 produced models exceeding the randomized baseline AUC, indicating that their corresponding knowledge graph derived gene networks contained predictive information beyond randomly selected genomic features. A comparison was performed with a similar, but closed source option in Claude Science. After asking it to generate 103 drug candidates with the same conditions, skills, and ability to query AlzKB, only 55 were above the randomized baseline. EcoXAI’s iterative hypotheses generation and its suite of diversified interactions with knowledge graphs are what separated the EcoXAI’s 79/103 (77%) to Claude Science’s 55/103 (53%). An interesting pattern emerged across successful EcoXAI hypotheses: direct drug targets alone frequently performed at or below the randomized baseline, whereas expanded biological interaction networks substantially improved predictive performance. This observation was reproduced across diverse drug classes including immune modulators, kinase inhibitors, metabolic drugs, calcium channel blockers, and GPCR-targeting compounds, suggesting that Alzheimer’s disease risk is better represented by wider networks than isolated molecular targets.

The final output of EcoXAI was a ranked list of 79 candidate drugs that exhibited improved predictive performance and biologically coherent mechanistic hypotheses derived entirely through autonomous reasoning over the biomedical knowledge graph. This demonstrates EcoXAI’s ability to move beyond automated model fitting by performing iterative scientific hypothesis generation, refinement, and biological interpretation with minimal human intervention.

### Case Study: Autonomous Discovery of Maraviroc

Among the 103 evaluated candidates, EcoXAI identified the CCR5 antagonist **Maraviroc** as one of the strongest repurposing hypotheses. Maraviroc is commonly used as an antiviral medication in treatment of HIV. Starting from experimentally validated drug targets, EcoXAI queried AlzKB to construct a 77-gene interaction network while explicitly excluding all Alzheimer’s disease-associated genes. On the 20% test split, this initial hypothesis achieved a cross-validated XGBoost AUC of 0.6013, exceeding the randomized baseline.

Without intervention and rather than stopping after obtaining a significant result, EcoXAI autonomously generated and evaluated refined hypotheses through additional knowledge graph expansion. Introducing second-order interaction “bridge” genes increased performance to an AUC of 0.6143, the highest-performing model was produced across all runs and comparable in performance to the best human curated models in prior studies (Orlenko et al., 2025). Feature importance analysis revealed that the predictive signal was driven primarily by downstream network genes rather than the direct drug target, suggesting that distributed biological interactions better capture Alzheimer’s disease susceptibility.

Although CCR5 was three hops removed from Alzheimer’s disease in the AlzKB graph and all AD-associated genes were explicitly excluded during hypothesis generation, subsequent literature review revealed substantial independent evidence supporting this connection. Previous studies have reported increased CCR5 expression in Alzheimer’s disease brains (Xia et al., 1998), implicated the CCL5/CCR5 axis in regulating amyloid-*β* and tau pathology through microglial signaling (Ma et al., 2023), and demonstrated that amyloid-*β* induces CCR5 expression in brain endothelial cells to promote immune cell trafficking across the blood-brain barrier (Li et al., 2009). EcoXAI independently determined Maraviroc as a drug candidate by hypothesis generation via agentic interaction with AlzKB and without knowledge of recent mouse model studies that found Maraviroc treating early-stage AD (Liu et al., 2026). Claude Science did not propose Maraviroc within its hypotheses. These findings suggest that EcoXAI recovered a biologically meaningful disease mechanism through autonomous knowledge graph reasoning rather than simply rediscovering known Alzheimer’s disease annotations.

## 5 Conclusion

EcoXAI represents a step toward autonomous AI-assisted biomedical data science by combining modern paradigms in agent harnesses, loop engineering, and unbound interaction with structured biomedical knowledge graphs. Unlike frameworks that act as solely a conversational assistant, EcoXAI proactively performs end-to-end scientific workflows including exploratory data analysis, knowledge retrieval, hypothesis generation, predictive modeling, and biological interpretation. Its modular, containerized architecture with harnessed agents and easily editable agent pipelines, enables reproducible deployment across diverse computational environments while allowing researchers to incorporate custom analytical tools, knowledge graphs, and local LLMs.

As agentic AI systems continue to mature, ensuring transparency, reproducibility, and interpretability are key especially in the biomedical domain. Like other autonomous AI systems, EcoXAI is susceptible to planning errors and cascading failures. To mitigate this, the framework enforces strict isolation across its pipeline stages, persisting intermediate code execution scripts and experimental logic so human researchers can inspect, reproduce, or modify tasks mid-loop.

Although demonstrated here for drug repurposing, the framework is readily extensible to other biomedical discovery tasks, including multi-omics integration, biomarker discovery, pathway analysis, and precision medicine applications. By coupling autonomous scientific reasoning with grounded biomedical knowledge, EcoXAI provides a scalable platform for accelerating hypothesis-driven biological discovery while maintaining the transparency required for biomedical research.

## Acknowledgment

This work is supported in part by funds from the Center for AI Research and Education at Cedars-Sinai Medical Center and grants from the National Institutes of Health USA (P30 AG094848, U01 AG066833, R01 LM014572, R01 LM010098, and K01 DA063751).

